# Distinct Brain Maturation Pathways Relate Environmental Variation to Cognitive Development

**DOI:** 10.64898/2026.04.04.716497

**Authors:** Jiadong Yan, Judy Chen, Bin Wan, Zhen-Qi Liu, Liang He, Paule Joanne Toussaint, Alan C. Evans, Sherif Karama

## Abstract

Adolescent cognitive ability develops within a complex landscape of environmental exposures, yet the patterns of brain maturation through which these exposures relate to cognitive development remain poorly understood. Resolving this requires an integrated longitudinal view of the multidimensional exposome, multimodal brain change, and cognitive growth. Here, we examined whether brain maturation mediates the relationship between the exposome and cognitive development in a large longitudinal cohort of 1,112 adolescents, using 4,882 multimodal brain features and 112 exposome variables spanning six domains. Brain maturation patterns associated with cognitive development showed moderate yet consistent spatial correspondence with those associated with the exposome, with the family environment domain showing the strongest correspondence. We identified four latent brain patterns associated with distinct components of the exposome-cognition relationship. Collectively, these results indicate that distinct exposome domains relate to cognitive development through distinct brain maturation pathways, which also exhibit different molecular annotation profiles.

## Introduction

Adolescent cognitive development is posited to be significantly shaped by the exposome, defined as the totality of environmental exposures (Tucker-Drob et al., 2013; Tucker-Drob & Briley, 2014), which encompasses multiple co-occurring exposure domains, including family context (Kwon et al., 2025; Wrulich et al., 2013), neighborhood environment (Fangfang et al., 2022; Song et al., 2024; Weuve et al., 2012), and early-life experiences (Chirwa & Yambayamba, 2026; Gennings et al., 2021; Yu et al., 2024). Among these, socioeconomic status, including family income, parental education, and housing quality, has consistently emerged as some of the most influential determinants of cognitive development (Kaplan et al., 2001; Korous et al., 2022; Tong et al., 2007). In addition, parenting practices and cognitive stimulation play critical roles in shaping individual differences in cognitive trajectories (Byford et al., 2012; Tucker-Drob & Harden, 2012). However, the mechanisms through which diverse exposures shape cognitive development remain poorly understood.

Understanding how the exposome influences cognitive development requires disentangling the role of brain maturation as a central neurodevelopmental pathway (Brito & Noble, 2014; Zhang et al., 2023; Zhi et al., 2024). During adolescence, the brain undergoes profound and regionally heterogeneous structural and functional changes (Blakemore & Choudhury, 2006; Giedd et al., 1999). Cortical gray matter undergoes region-specific thinning from primary sensorimotor areas to higher-order association cortices (Gogtay et al., 2004; Tamnes et al., 2017), while white matter microstructure matures progressively to support the development of long-range structural connectivity (Lebel & Beaulieu, 2011). In parallel, functional network organization shifts toward greater between-network segregation and within-network integration (Fair et al., 2009; Luna et al., 2015). These maturational processes both increase cognitive ability (Kail & Salthouse, 1994; Luna et al., 2004) and render a period of heightened susceptibility to environmental exposures (Blakemore, 2012; Paus et al., 2008).

Past literature has established bilateral relationships from exposome or cognitive ability and brain maturation. Socioeconomic disadvantage is associated with reduced cortical surface area in frontal and temporal regions (Hair et al., 2015; Noble et al., 2015), while environmental adversity appears to alter connectivity within default mode and frontoparietal networks (Ellwood-Lowe et al., 2021; Tooley et al., 2021). Likewise, variation in cortical surface area or connectivity correlates with differences in cognitive performance (Finn et al., 2015; Khundrakpam et al., 2017). Yet whether brain maturation integrates these bilateral links into a coherent developmental pathway, and whether different exposome domains map onto cognitive development through shared or distinct neural substrates, remains unresolved. One reason is that few studies have modeled developmental changes across brain and cognition simultaneously (Farah, 2017). Without concurrent longitudinal measurement of both, observed associations may reflect stable individual differences rather than ongoing maturational processes. Another is that prior work has typically examined narrow subsets of either brain measures or environmental variables, limiting integrated understanding of how distinct exposome domains relate to cognitive development through the brain (Brito & Noble, 2014; Zhang et al., 2023; Zhi et al., 2024).

In this study, we present a comprehensive, longitudinal framework using the Adolescent Brain Cognitive Development (ABCD) Study (*N* = 1,112; age 9-15 years; Casey et al., 2018) to investigate the relationship between exposome domains and neurocognitive development. We characterized the relationships among 112 exposome variables spanning six domains, 4,882 multimodal brain features derived from structural and functional MRI, and general cognitive ability (*g*), a latent factor capturing shared variance across cognitive domains (Barbey, 2018; Colom et al., 2010). The overview of the analytical workflow was shown in **Fig. 1**. We discovered that brain maturation associated with *g* development tended to also be sensitive to the exposome. Beyond this convergence, mediation analyses revealed that different exposome domains relate to *g* development through distinct neural substrates. Our collective findings demonstrate that while the exposome and *g* converge on similar patterns of brain maturation at the aggregate level, different exposome domains link to cognitive growth through distinct candidate neural pathways.

**Fig. 1.**
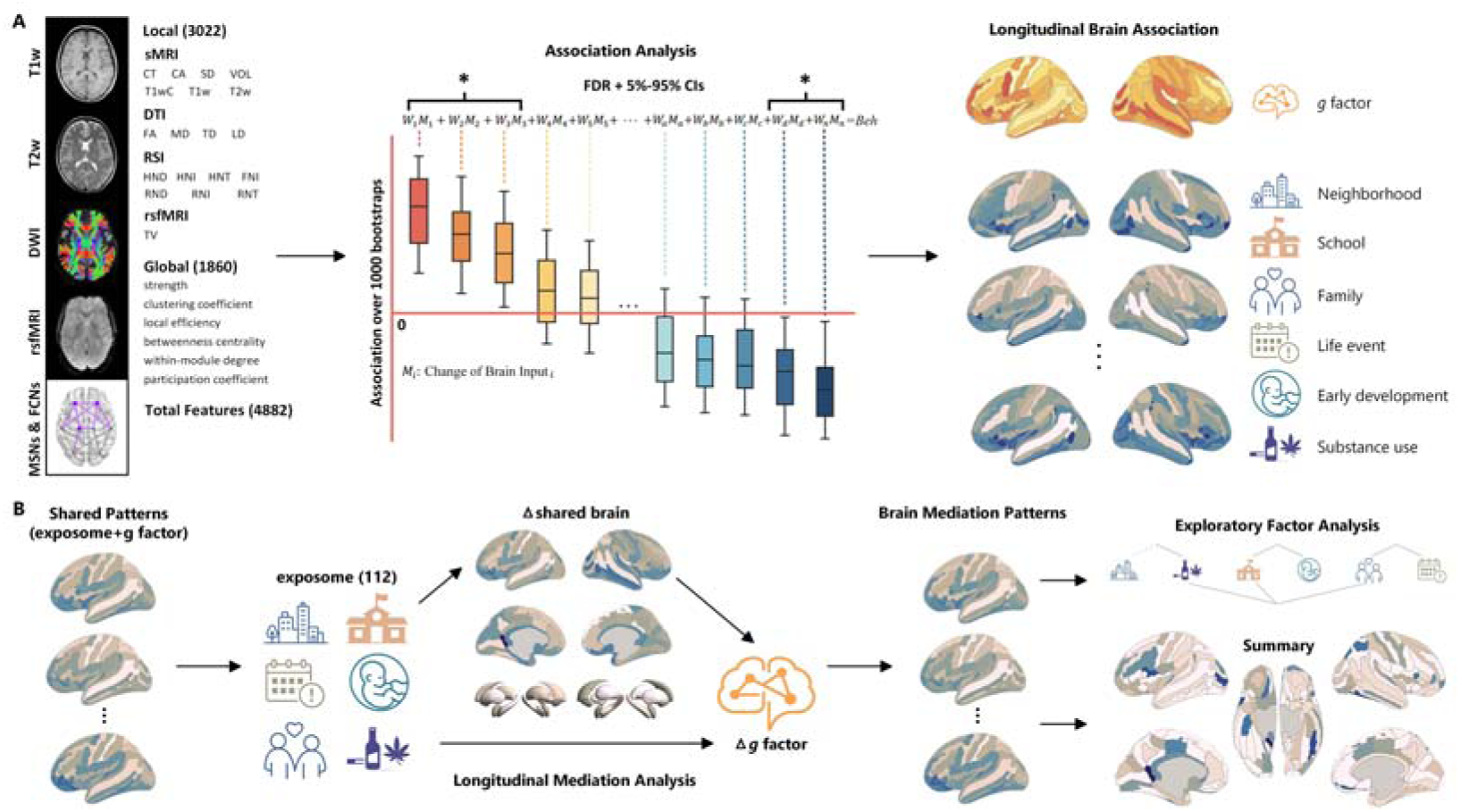
Overview of the analytical workflow. (A) A total of 4,882 local and global brain features were extracted from multimodal neuroimaging data (sMRI, DTI, RSI, and rsfMRI). Longitudinal associations between brain and both the *g* factor and 112 exposome variables were subsequently examined. (B) Shared brain patterns jointly associated with the *g* factor and the exposome were first identified. Longitudinal mediation analyses were then conducted to determine which brain changes mediated the effects of exposome on *g* development. The resulting brain mediation patterns were further characterized through exploratory factor analysis to identify distinct factors underlying brain-mediated effects. A summary pattern was also derived to provide an integrated representation of the overall mediation effects.

## Results

### Brain maturation is longitudinally associated with cognitive development and the exposome

Longitudinal association analyses revealed that brain maturation was broadly associated with *g* development, with the strongest associations concentrated in frontal, temporal, and occipital regions (**Fig. 2A**). For exposome domains, the three lobes with the highest association strength were consistently the temporal, limbic, and insular lobes across five of the six domains; the family environment domain was an exception, with the frontal lobe replacing the insula among the top three (**Fig. 2B**). Across all seven patterns, the temporal lobe contributed the largest proportional share of association strength, while the parietal lobe and subcortex contributed the least (unpaired t-tests, *p* < 0.05; **Fig. 2C**).

**Fig. 2.**
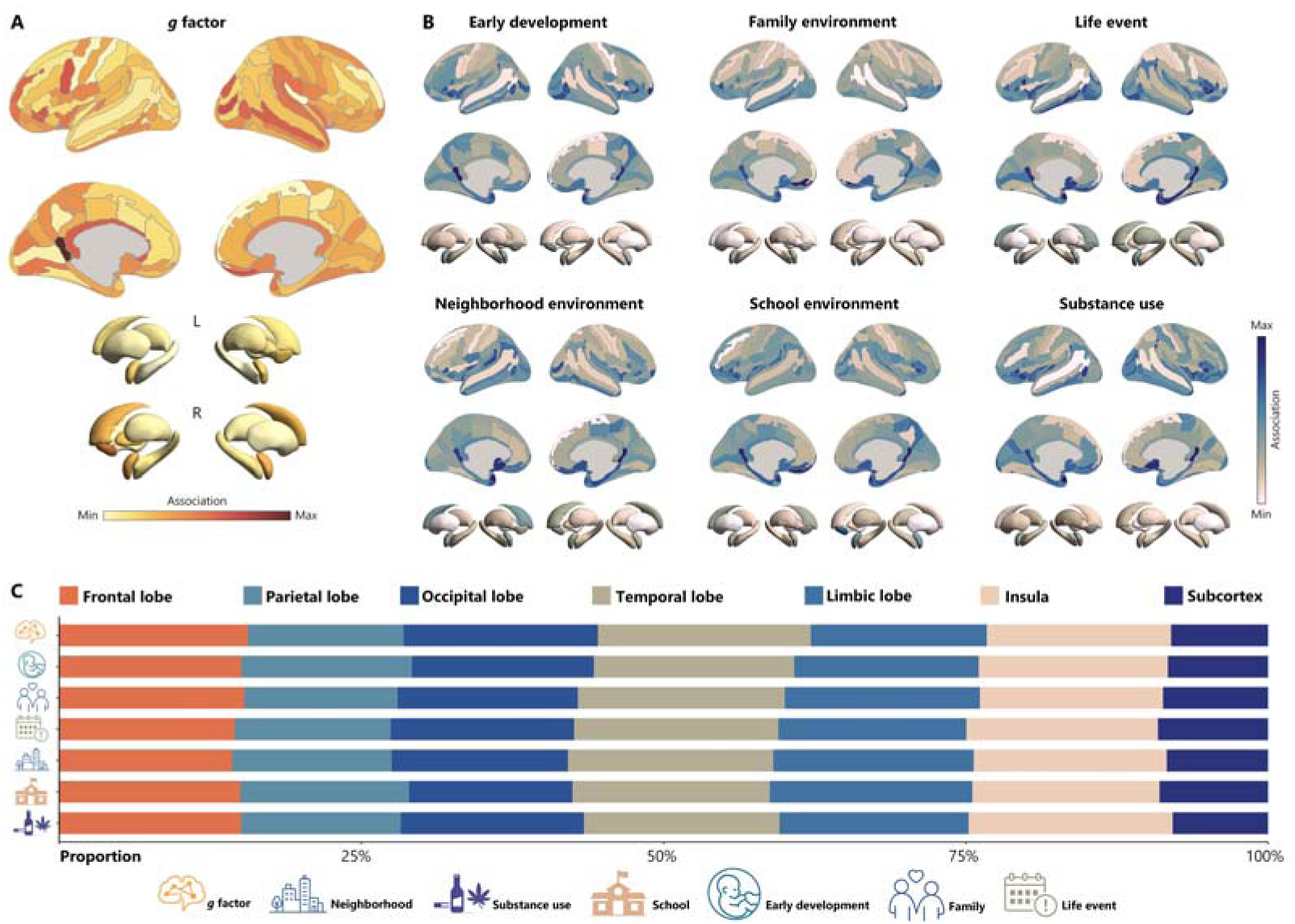
Longitudinal brain association patterns for *g* development and the exposome. (A) Brain association pattern for *g* development. Color intensity reflects the magnitude of association, normalized to a unit sum to facilitate cross-region comparison. (B) Domain-averaged brain association patterns for the six exposome domains. Color intensity reflects the magnitude of association, normalized to a unit sum within each domain to facilitate cross-region comparison. (C) Lobe-level composition of brain association patterns for *g* development and the six exposome domains, expressed as each lobe’s proportional share of the summed association strength within each pattern.

### Brain maturation patterns are broadly shared between cognition and the exposome

Similarity analyses revealed moderate correspondence between the *g* association pattern and exposome association patterns, with cosine similarities ranging from 0.20 to 0.34 across the 112 exposome measures. Among exposome measures, the top 10 were predominantly related to parental mental health and socioeconomic dimensions of the neighborhood and school environment (**Fig. 3A**). At the domain level, family environment showed the highest similarity to the *g*-related brain maturation pattern, significantly exceeding early development, life events, and substance use domains (unpaired t-tests, *p* < 0.05).

**Fig. 3.**
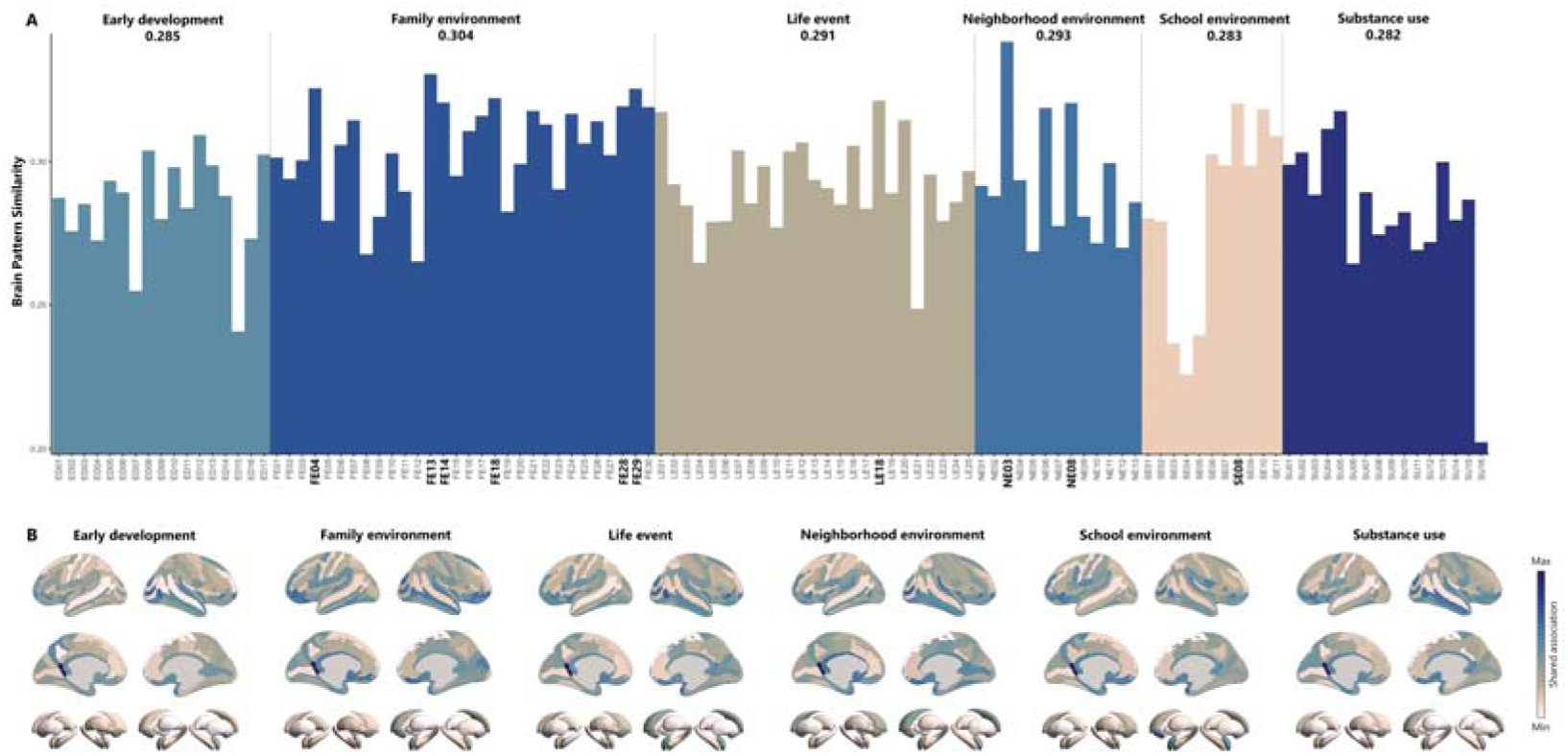
Shared longitudinal brain association patterns between *g* development and the exposome. (A) Brain pattern similarity between the longitudinal association patterns of *g* development and each of the 112 exposome measures, grouped by domain. The number above each domain indicates the domain-averaged similarity value. Bold labels denote the ten exposome measures with the highest similarity to the *g* association pattern: area deprivation index (NE03), parental stress-related perseveration (FE13), marital status (FE04), parental DSM-5 hyperactivity (FE29), parental attention problems (FE18), sleep arousal (LE18), parental anxiety/depression (FE14), noise exposure (NE08), mental health and substance use service access (SE08), and parental DSM-5 inattention (FE28). ED, FE, LE, NE, SE, and SU denote early development, family environment, life events, neighborhood environment, school environment, and substance use, respectively. (B) Domain-averaged shared brain association patterns between *g* development and each exposome domain. Color intensity reflects the magnitude of shared association, normalized to a unit sum within each domain to facilitate cross-region comparison.

We further examined shared longitudinal brain association patterns between *g* and each of the six exposome domains. The six domain-level shared patterns were highly convergent. The most strongly shared regions encompassed the posterior-ventral cingulate cortex, primary and associative auditory cortex, higher-order visual cortex, orbitofrontal and olfactory cortex, and the anterior transverse collateral sulcus (**Fig. 3B**). Specific ROI names are provided in **Supplementary Note 1**.

### Brain maturation links exposome to cognitive development

Mediation analyses across all six exposome domains identified distributed brain regions through which brain maturation mediates associations between the exposome and *g* development. The summary mediation pattern, aggregated across all 112 exposome measures, implicated a broad set of cortical and subcortical regions (**Fig. 4A**). Prominent regions included the posterior-ventral cingulate cortex, auditory cortex, higher-order visual cortex, orbitofrontal cortex, and parietal cortex. Specific ROI names are provided in **Supplementary Note 2**.

**Fig. 4.**
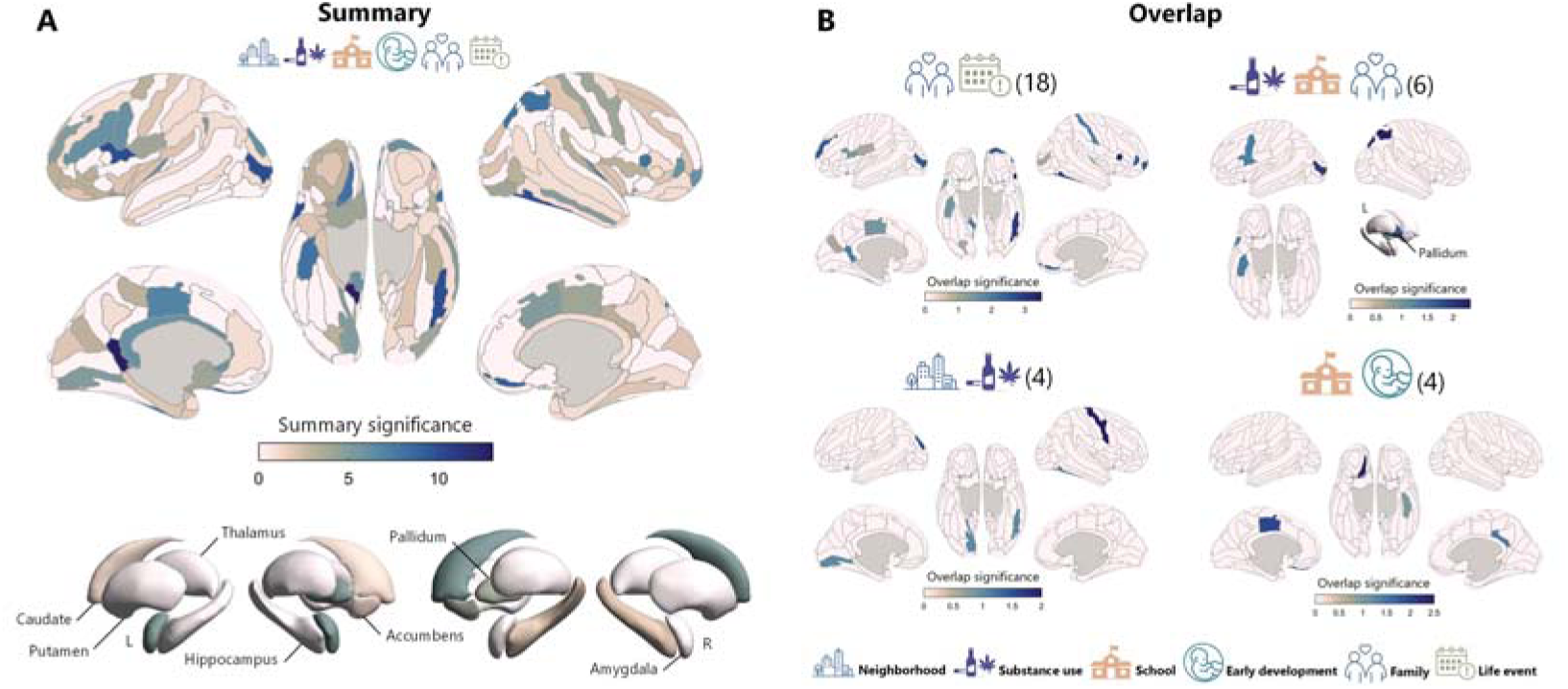
Brain mediation patterns linking exposome measures to *g* development. (A) Summary brain mediation pattern aggregated across all 112 exposome measures. Color intensity reflects the total number of significant mediation effects across exposome measures within each brain region. (B) Factor-specific consensus brain mediation patterns derived from EFA of domain-level mediation patterns. Each panel corresponds to one latent factor, with the number of significant brain regions indicated in parentheses. Only brain regions showing significant mediation effects across all exposome domains within a given factor were retained, and color intensity reflects the average mediation count across those domains. Exposome icons above each panel identify the exposome domains contributing to that brain pattern.

To identify consensus mediation patterns across exposome domains, EFA-based method was applied to domain-level mediation patterns, yielding four latent factors (**Fig. 4B** and **Supplementary Fig. 1**). Factor 1, loading on family environment and life events, showed the most extensive mediation footprint (18 regions), encompassing cingulate cortex, orbitofrontal cortex, opercular inferior frontal gyrus, higher-order visual cortex, lateral occipito-temporal cortex, and additional frontal, parietal, and perisylvian regions. Factor 2, loading on family environment, substance use, and school environment, implicated parietal cortex, higher-order visual cortex, pallidum, opercular inferior frontal gyrus, inferior precentral sulcus, and anterior transverse collateral sulcus (6 regions). Factor 3, loading on neighborhood environment and substance use, involved precentral gyrus, occipital cortex, lingual gyrus, and lateral occipito-temporal cortex (4 regions). Factor 4, loading on school environment and early development, was centered on orbitofrontal cortex, cingulate cortex, and anterior transverse collateral sulcus (4 regions). Specific ROI names are provided in **Supplementary Note 3**. The implicated regions showed limited overlap across factors, suggesting that different exposome domains engage distinct spatial patterns of brain maturation in shaping *g* development. Results of multiple comparisons are provided in **Supplementary Fig. 2** and **Supplementary Note 4**.

### Distinct molecular profiles underline the four brain mediation patterns

To investigate the molecular profiles associated with the previous four patterns, we tested their spatial correspondence with neuromaps (Hansen et al., 2022; Markello et al., 2022). Factor 1 (related to family environment and life events) was positively correlated with resting cerebral blood flow (*r* = 0.15). Factor 2 (related to family environment, school environment, and substance use) showed no significant correspondence with any annotation map. Factor 3 (related to neighborhood environment and substance use) was negatively correlated with PET-derived maps of 5-HT_6_ serotonin receptor (*r* = -0.19), mGluR_5_ (*r* = -0.18), SV2A synaptic density (*r* = -0.19), GABA_A_ receptor (*r* = -0.18), GABA_A_ α_5_ subunit (*r* = -0.19), and M1 muscarinic acetylcholine receptor (*r* = -0.17). Factor 4 (related to early development and school environment) was negatively correlated with PET-derived maps of 5-HT_1B_ serotonin receptor (*r* = -0.18), mGluR_5_ (*r* = -0.15), and α_4_β_2_ nicotinic acetylcholine receptor (*r* = -0.21). All reported correlations survived spin tests (*p* < 0.05).

## Discussion

To investigate how the exposome relates to adolescent cognitive development through brain maturation, we conducted large-scale analyses of extensive brain, exposome, and cognitive measures to map their relationships. Extending previous work (Zhang et al., 2023; Zhi et al., 2024), we not only investigated comprehensive brain maturation patterns that mediate exposome-*g* associations but also identified distinct exposome domains whose associations with *g* development are carried by different neural substrates.

### Shared brain maturation patterns between cognition and the exposome

The shared components of the *g*- and exposome-related brain association patterns were similar across the six exposome domains. Notably, regions contributing to these patterns mainly included the posterior-ventral cingulate cortex, orbitofrontal cortex, and higher-order auditory and visual association areas, overlapping with networks described in the Parieto-Frontal Integration Theory (P-FIT) network proposed to underlie general cognitive ability (Basten et al., 2015; Colom et al., 2006; Jung & Haier, 2007). This suggests that the cortical systems associated with *g* overlapped with those related to environmental variation. Comparatively, primary sensory cortices contributed less to the shared components than association cortices. This pattern is consistent with differences in developmental timing, with primary sensory regions largely established earlier in life and association cortices continuing to mature through adolescence (Gogtay et al., 2004; Shaw et al., 2008; Tooley et al., 2021), creating a longer window of environmental susceptibility. This maturational gradient aligns with hierarchical models of cortical organization, in which a principal gradient from sensorimotor to transmodal cortex has been described (Margulies et al., 2016). Regions toward the association end of this gradient continue to mature later and may be more sensitive to environmental variation (Sydnor et al., 2021; Valk et al., 2022).

### Family environment as a prominent source of environmental variation

Among exposome factors, family environment showed the highest similarity to the *g*-related brain maturation pattern. This finding aligns with twin studies that demonstrate the substantial contribution of shared environment to childhood cognitive ability (Tucker-Drob & Harden, 2012). Within this domain, parental mental health emerged as the most significant factor, consistent with earlier findings linking parental psychopathology to offspring neurocognitive development (Goodman et al., 2011; Goodman & Gotlib, 1999). Further neuroimaging studies have illustrated strong associations between parental psychopathology and reduced cortical thickness and surface area in frontal and temporal regions in children (Lebel et al., 2016; Xerxa et al., 2024). These results highlight the family exposures as a key source of environmental variation to systematically account for when studying brain maturation and cognitive growth.

It is worth noting that family environment measures such as parental mental health and socioeconomic status are themselves partly heritable, as parental genotypes shape both the environment they provide and the genes they transmit, which is known as passive gene-environment correlation (Plomin et al., 1985; Scarr & McCartney, 1983). The associations we observe therefore likely reflect intertwined environmental, social, and genetic processes rather than purely environmental influences. Critically, our findings remain meaningful regardless of these mixed origins.

### Distributed brain mediators of exposome-cognition associations

Mediation analyses revealed that the brain regions through which maturation mediates exposome-cognition associations are spatially distributed, primarily implicating the posterior cingulate, orbitofrontal, auditory, higher-order visual, and intraparietal cortices. Here, the posterior cingulate cortex (PCC) emerged as a prominent mediator of the associations between brain maturation and *g* development, consistent with previous findings of its role as a core hub supporting cognition and memory consolidation, and its sensitivity to environmental variation during maturation (Gogtay et al., 2004; Leech & Sharp, 2014; Raichle, 2015). Furthermore, its integral position within the default mode network suggests that the PCC may represent a central hub through which exposome-related developmental variation becomes associated with cognitive development (Gong et al., 2021; Pozzi et al., 2021; Sherman et al., 2014; Whitfield-Gabrieli et al., 2020). Although sensory cortices contributed less to the previous shared *g*-exposome brain maturation patterns, some regions still emerged as significant mediators of the exposome-*g* association. This may reflect a bottleneck at where sensory cortices perform the initial processing and where environmental information enters the cortical hierarchy; variation in their maturational integrity may constrain downstream cognitive operations regardless of the efficiency of higher-order association regions (Murray et al., 2014).

### Distinct brain maturation patterns linking exposome domains to cognitive development

While similarity analyses revealed a common brain association pattern between *g* and the exposome domains, our mediation analyses showed that different exposome domains implicate non-overlapping sets of mediating brain regions. The first factor group comprising of family environment and life events showed the most extensive mediation footprint, spanning cingulate, orbitofrontal, opercular, visual, and parietal cortices. These regions collectively support core components of general cognitive ability (Deary et al., 2010; Jung & Haier, 2007), and likely reflect the pervasive and cumulative nature of daily family life (Hackman et al., 2010; Luby et al., 2012). Life events loaded onto the same factor as family environment, consistent with ecological models demonstrating that stressful events are typically experienced within and filtered through the family context (Bronfenbrenner & Morris, 2007). This suggests that life events may be linked to cognitive outcomes through the same cortical systems that mediate the associations between family environment and *g* development.

The second factor group includes family environment, substance use, and school environment, and implicates the parietal cortex, pallidum, and opercular regions. In our cohort’s age range (9-15 years), substance use measures capture mainly household rules, parental attitudes toward use, and perceived availability, rather than actual use or abuse. Taken together with school and family environment, this factor may capture structured regulatory contexts associated with cognitive development through behavioral regulation. Moreover, the involvement of the pallidum and the intraparietal sulcus, key players in reward and attentional control respectively (Casey et al., 2008; Corbetta & Shulman, 2002; Galvan, 2010; Smith et al., 2009; Steinberg, 2008), suggests that regulatory environments may be linked to cognition through the coordinated maturation of reward-driven and attentional systems.

Our third factor combining neighborhood environment and substance use was centered on motor and visual cortices. We posit that sensorimotor cortices, the gateway of initial environmental processing information, may be the most sensitive to restrictive sensorimotor experiences during adolescent development in disadvantaged neighborhoods that are often associated with higher crime, substance use, and reduced access to safe environments for physical play and exploration (Gao et al., 2015; Leventhal & Brooks-Gunn, 2000; Rakesh et al., 2021). Consequently, our findings suggest that environmentally constrained sensorimotor cortical maturation may possibly limit higher-order cognitive processing and ability. Finally, the fourth factor loaded on early development and school environment. Both domains are highly tied to cognitive functioning itself (Gennings et al., 2021; Wrulich et al., 2013), and the shared brain pattern resembled the pattern observed in our previous work on brain-cognition associations (Yan et al., 2026), further suggesting it may capture neural correlates of cognitive ability.

Overall, the limited spatial overlap across factors suggests that different exposome domains are linked to cognitive development through largely distinct neural substrates, and that understanding how the exposome relates to cognitive development requires accounting for domain-specific brain patterns (McLaughlin & Sheridan, 2016). Taken together with the convergent *g*-exposome brain patterns described earlier, these findings suggest that while general cognitive ability reflects a common neurocognitive architecture (Johnson et al., 2008), the developmental pathways through which environmental variation relates to that architecture may be partially dissociable.

### Molecular underpinnings of the distinct brain mediation patterns

The molecular annotations help differentiate the previous four mediation patterns. Factor 1 (family environment and life events) correlated positively with resting cerebral blood flow (Satterthwaite et al., 2014), which localizes this factor to metabolically demanding association cortex. This suggests that chronic family exposures may therefore relate to *g* development through cortical systems with high baseline metabolic activity. Factor 2 (family environment, school environment, and substance use) showed no correspondence with any molecular map. This is consistent with our earlier observation that Factor 2’s brain pattern depends on coordinated maturation of the pallidum and intraparietal regions, which are not characterized by a single neurochemical system. Both regions have been implicated in reward processing (Haber & Knutson, 2009) and attentional control (Corbetta & Shulman, 2002). Factor 3 (neighborhood environment and substance use) correlated negatively with several receptor and synaptic density maps (5-HT_6_, mGluR_5_, GABA_A_, GABA_A_ α_5_, M1 muscarinic, and SV2A) that are concentrated in transmodal association cortex (Hansen et al., 2022; Sydnor et al., 2021). These findings indicate that Factor 3 sits toward the sensorimotor pole, which is dominated by motor and visual cortex. Neighborhood disadvantage may thus shape *g* by constraining sensorimotor maturation. Factor 4 (early development and school environment) correlated negatively with 5-HT_1B_, mGluR_5_, and α_4_β_2_ nicotinic receptor maps. These receptors are involved in synaptic plasticity and cholinergic modulation of cortical activity (Picciotto et al., 2012). Overall, the four factors aligned with neurochemically distinct brain systems, suggesting that different exposome domains relate to cognitive development through different molecular profiles.

## Limitations

Several limitations of the current study should be noted. First, despite the longitudinal design, our mediation analyses remain statistical rather than causal. Therefore, our findings should be interpreted as identifying brain maturation patterns consistent with a mediating role, rather than establishing causal pathways. Causal inference would require experimental or quasi-experimental approaches. Second, all exposome measures used were assessed at baseline, so changes in environmental conditions over the follow-up period were not incorporated; future work incorporating time-varying exposome measures would capture a more dynamic illustration of exposome-brain-cognition relationships. Third, we did not directly model genomic contributions. As discussed above, many exposome measures, particularly those related to the family environment, are partly heritable and subject to intergenerational continuity (Kendler & Baker, 2007; Plomin & Bergeman, 1991). Future work incorporating genetically informed designs would help formally partition environmental and genetic contributions to exposome-brain-cognition associations (Cheesman et al., 2020; Kong et al., 2018; Nivard et al., 2024).

## Conclusions

This study provides a systematic characterization of brain maturation patterns associated with the exposome and adolescent cognitive development. At the aggregate level, exposure factors and cognition converge on overlapping brain regions, with family environment, particularly parental mental health, showing the strongest correspondence with brain maturation associated with cognitive development. Beneath this convergence, however, four distinct brain maturation patterns, each with a different molecular signature, mediate the relationship between specific exposome domains and cognitive growth. Together, these findings suggest that environmental variation relates to adolescent cognitive development through multiple partially dissociable neurodevelopmental pathways embedded within the general cognitive architecture.

## Methods

### Participants

The ABCD 5.1 release cohort includes 11,868 participants recruited across 22 sites (Casey et al., 2018). After quality control, (Hagler et al., 2019) the final longitudinal sample consisted of 1,112 participants (553 females; age range: 9-15 years). Full details are provided in **Supplementary Table 1** and **Supplementary Note 5**. All procedures were approved by the Institutional Review Board at the University of California, San Diego, and written informed consent was obtained from all participants.

### Brain features

#### Brain structure

Multimodal structural brain measures were derived from T1-weighted (T1w), T2-weighted (T2w), and diffusion MRI (dMRI) data processed using the ABCD Study pipeline (Casey et al., 2018; Hagler et al., 2019; **Fig. 1**). Cortical and subcortical regions were delineated using the Destrieux atlas (148 ROIs) and the ASEG atlas (14 ROIs), respectively (Destrieux et al., 2010; Fischl et al., 2002). For each cortical ROI, 18 local structural measures were extracted across three modalities: seven sMRI metrics, four DTI metrics, and seven RSI metrics (White et al., 2014). Subcortical ROIs yielded 14 measures per region, with the four cortical-specific metrics excluded. Collectively, these measures produced 2,860 local structural features (148×18 + 14×14). Morphometric similarity networks (MSNs, Seidlitz et al., 2018) were subsequently constructed from these local features, and six graph-theoretic measures were computed per ROI (Rubinov and Sporns, 2010), yielding an additional 972 global structural features (162×6). Detailed descriptions of all measures are provided in **Supplementary Note 6**.

#### Brain function

Functional MRI data were preprocessed using fMRIPrep (Esteban et al., 2018, 2020) and denoised using XCP-D (Mehta et al., 2024). Following preprocessing, functional time series were projected onto the fsLR surface space for subsequent analyses. Using the same cortical parcellation scheme, local functional measures for each of the 162 ROIs were quantified as the temporal variance of the BOLD signal. Functional connectivity networks were then constructed (Biswal et al., 1995), and six graph-theoretic measures were computed per ROI, yielding 888 global functional features (148×6). Detailed preprocessing procedures are described in **Supplementary Note 7**.

In total, 4,882 brain measures were extracted, comprising 2,860 local structural, 972 global structural, 162 local functional, and 888 global functional features.

### Exposome measures

A total of 112 exposome measures were included, spanning six domains that reflect influences across early development, family environment, life events, neighborhood environment, school environment, and substance use (Modabbernia et al., 2021; Zhi et al., 2024). The early development domain comprised 17 measures capturing prenatal, perinatal, and early postnatal factors. The family environment domain comprised 30 measures indexing household structure, socioeconomic context, caregiving characteristics, and psychosocial functioning. The life events domain comprised 25 measures reflecting social experiences, daily activities, health-related behaviors, and stress exposure. The neighborhood environment domain comprised 13 measures characterizing the physical, environmental, and socioeconomic attributes of the residential context. The school environment domain comprised 11 measures assessing academic performance, educational support, disciplinary experiences, and school engagement. Finally, the substance use domain comprised 16 measures capturing substance-related expectancies, perceived availability, parental permissiveness, and early experimentation. Full details of all 112 measures are provided in **Supplementary Table 2**.

### Cognitive assessment

Cognitive ability was assessed using six subtests drawn from the ABCD dataset, selected to represent multiple cognitive domains (Yan et al., 2025, 2026): vocabulary, attention, reading, processing speed, episodic memory, and visuospatial accuracy (Luciana et al., 2018). Of the original 12 subtests in the ABCD battery, those with incomplete data attributable to COVID-19 disruptions were excluded, yielding the six subtests used here. A principal component analysis (PCA) was then conducted on the six subtest scores, and the first principal component was retained as an index of *g*, consistent with established practices in the field (Barbey, 2018; Colom et al., 2010). The *g* derived from these six subtests was highly correlated with that derived from the complete battery (*r* > 0.85), supporting the representativeness of our measure.

### Longitudinal association analysis

We applied an association model (Yan et al., 2025, 2026; Zhou et al., 2025) to examine longitudinal associations of changes in 4,882 brain features with 113 phenotypes (112 baseline exposome measures and 1 *g* change). For each phenotype, we implemented a multivariate regression model in which all brain features were entered simultaneously, allowing interdependencies among them to be accounted for (**Fig. 1A**). This framework is identical to that used in a previous study (Yan et al., 2026), in which multicollinearity and coefficient stability were systematically evaluated and shown to be adequately controlled. All models controlled for age, total brain volume, sex, handedness, ethnicity, study site, family, and principal components of genetic ancestry (Price et al., 2006). Given 113 phenotypes, 113 separate models were fitted. A brain feature was considered significantly associated with a given phenotype only if it satisfied two criteria (Yan et al., 2026): (1) its mean regression coefficient across bootstrap iterations differed from zero after Bonferroni correction, and (2) its 5-95% bootstrap confidence interval did not cross zero. For significant features, association strength was defined as the absolute value of the mean coefficient across bootstrap iterations; non-significant features were assigned a value of zero. Additional technical details are provided in **Supplementary Note 8**.

Since our analysis adopted the full set of comprehensive brain features, comparing effects across regions required a common summary. We therefore obtained the overall association strength of each brain region by summing the absolute association values across all features within that region. After obtaining exposome-specific longitudinal brain association patterns, we averaged patterns across exposome measures within each of the six domains to derive domain-level brain association patterns. We also computed the cosine similarity between the *g* association pattern and each of the 112 exposome association patterns to characterize their spatial alignment. For each exposome measure, the shared brain association pattern was further quantified by averaging association values across features significantly associated with both *g* development and that exposome measure, which were subsequently used for mediation analyses.

### Longitudinal mediation analysis

We conducted longitudinal mediation analyses to identify brain maturation patterns that statistically mediate the associations between baseline exposome and *g* development (**Fig. 1B**). A nonparametric bootstrap approach was used to evaluate mediation significance (Alfons et al., 2022), with 1,000 bootstrap iterations applied to all models. All mediation analyses were conducted within the same covariate framework as the primary association analyses. For each exposome measure, multiple comparisons across all tested shared brain mediators were corrected using Bonferroni adjustment. The brain mediation pattern for each exposome measure was represented as the total number of significant mediation effects summed across all features within each brain region. Additional technical details are provided in **Supplementary Note 9**.

After obtaining exposome-specific brain mediation patterns, we summed patterns within each of the six exposome domains to derive domain-level brain mediation patterns. We then conducted an exploratory factor analysis (EFA) on the domain-level mediation patterns to identify latent factors that group exposome domains sharing common brain-mediated effects on *g* development (Johnson et al., 2008). EFA hyperparameters were set following established practice (Comrey & Lee, 2013; Johnson et al., 2008), with technical details reported in **Supplementary Note 9**. Within each identified factor, a consensus mediation pattern was derived by retaining brain regions present in all constituent exposome domain patterns, with the counts of the retained brain regions averaged across domains. Finally, we summed mediation patterns across all 112 exposome measures to derive a summary pattern reflecting the overall brain mediation effect.

### Annotation enrichment analysis

To contextualize the latent brain patterns identified by EFA, we used the neuromaps toolbox (Hansen et al., 2022; Markello et al., 2022) to test their spatial correspondence with reference brain maps spanning metabolism, neurotransmitter receptors and transporters, and synaptic density. Reference maps were resampled to fsaverage space and parcellated using the Destrieux atlas to match the resolution of the latent patterns. Pearson correlations were assessed against a spatial null distribution (spin test; Markello & Misic, 2021).

## Data and code availability

The data used in this study were obtained from the ABCD Study, release 5.1, under the terms of a data use agreement. Raw data cannot be shared directly by the authors but are accessible through application to the NIMH Data Archive (NDA). Researchers with approved access can use the provided scripts to reproduce the analyses. All analysis code is accessible online: https://github.com/JDYan/Longitudinal-Mediation.

## Acknowledgements

Sherif Karama is supported by the Canadian Institutes of Health Research. Alan Evans is supported by CFREF/HBHL. Jiadong Yan is supported by the China Scholarship Council. Zhen-Qi Liu is supported by the Newton International Fellowship, Royal Society, UK. Data used in this study were obtained from the ABCD Study (https://abcdstudy.org), held in the NDA. The ABCD Study is a multisite, longitudinal study designed to recruit over 10000 children aged 9-10 and follow them for 10 years. A complete list of participating sites and investigators is available at https://abcdstudy.org/consortium_members/.

